# Loss of function of the mitochondrial peptidase PITRM1 induces proteotoxic stress and Alzheimer’s disease-like pathology in human cerebral organoids

**DOI:** 10.1101/2020.01.27.919522

**Authors:** Dina Ivanyuk, María José Pérez, Vasiliki Panagiotakopoulou, Gabriele Di Napoli, Dario Brunetti, Rawaa Al-Shaana, Stephan A. Kaeser, Mathias Jucker, Massimo Zeviani, Carlo Viscomi, Michela Deleidi

**Affiliations:** German Center for Neurodegenerative Diseases (DZNE), Tübingen, Germany; Department of Neurodegenerative Diseases, Hertie Institute for Clinical Brain Research, University of Tübingen, Tübingen, Germany; MRC-Mitochondrial Biology Unit, Cambridge CB2 0XY, UK; Department of Cellular Neurology, Hertie Institute for Clinical Brain Research, University of Tübingen, Tübingen, Germany

**Author notes:** These authors contributed equally to the work (Dina Ivanyuk, María José Pérez). Manuscript correspondence: Michela Deleidi, MD, PhD German Center for Neurodegenerative Diseases (DZNE) Tübingen within the Helmholtz Association, Department of Neurodegenerative Diseases, University of Tübingen Otfried-Müller. Str 23 72076 Tübingen-Germany.

## Abstract

Mutations in pitrilysin metallopeptidase 1 (*PITRM1*), a mitochondrial protease involved in mitochondrial precursor processing and degradation, result in a slow-progressive syndrome, characterized by cerebellar ataxia, psychotic episodes and obsessive behavior as well as cognitive decline. To investigate the pathogenetic mechanisms of mitochondrial presequence processing, we employed cortical neurons and cerebral organoids generated from PITRM1 knockout human induced pluripotent stem cells (iPSCs). PITRM1 deficiency strongly induced mitochondrial unfolded protein response (UPR^mt^) and enhanced mitochondrial clearance in iPSC-derived neurons. Furthermore, we observed increased levels of amyloid precursor protein and amyloid β in PITRM1 knockout neurons. However, neither cell death nor protein aggregates were observed in 2D iPSC-derived cortical neuronal cultures. On the contrary, cerebral organoids generated from PITRM1 knockout iPSCs spontaneously developed over time pathological features of Alzheimer’s disease (AD), including accumulation of protein aggregates, tau pathology, and neuronal cell death. Importantly, we provide evidence for a protective role of UPR^mt^ and mitochondrial clearance against impaired mitochondrial presequence processing and proteotoxic stress. In summary, we propose a novel concept of PITRM1-linked neurological syndrome whereby defects of mitochondrial presequence processing induce an early activation of UPR^mt^ that, in turn, modulates cytosolic quality control pathways. Thus our work supports a mechanistic link between mitochondrial function and common neurodegenerative proteinopathies.

## Introduction

Mitochondrial dysfunction has been described as a common hallmark of neurological diseases ^1^. However, mitochondria are often considered to be a secondary target, rather than the actual disease driver in these conditions. We have recently described three independent families carrying missense loss of function mutations in *pitrilysin metallopeptidase 1* (PITRM1), resulting in an age-dependent, progressive, neurological syndrome ^2, 3^. Patients suffer from progressive cerebellar dysfunction leading to cerebellar atrophy, and psychiatric manifestations including obsessive behavior, anger attacks, and psychosis ^2, 3^. Interestingly, some of these patients showed a deterioration of their cognitive functions with a slow progression till their late sixties ^2, 3^. Human PITRM1, also known as presequence peptidase (hPreP), is a nuclear-encoded mitochondrial gene, expressed in a number of tissues, including muscles and different brain regions, e.g. cortex, hippocampus, cerebellum, and tectum ^4^. PITRM1 was initially identified in *Arabidopsis thaliana* as a protease that degrades targeting peptides in mitochondria and chloroplasts ^5^. Human PITRM1 is a mitochondrial matrix enzyme that digests the mitochondrial-targeting sequences (MTS) of proteins imported across the inner mitochondrial membrane, after their cleavage from protein precursors by the mitochondrial matrix presequence peptidase (MPP). When the MTS are not properly degraded, they accumulate within the mitochondrial matrix causing dissipation of the mitochondrial membrane potential and mitochondrial dysfunction ^6, 7^. The incomplete processing of mitochondrial preproteins leads to their destabilization, resulting in alterations of mitochondrial proteostasis ^8^. *In vitro* studies using recombinant PITRM1 have shown that, besides MTS, the enzyme is also involved in the degradation of short, unstructured peptides and amyloid beta (Aβ) peptides ^8-11^. Interestingly, Aβ peptides inhibit the activity of CYM1, the PITRM1 orthologue in yeast, leading to accumulation of precursor proteins ^8^. Experimental work in mouse models supports a causal role for PITRM1 in neurodegenerative dementia, whereby loss of PITRM1 function leads to a progressive, neurodegenerative phenotype characterized by hindlimb clasping, impairment in motor coordination, and basal-ganglia related movement control ^2^. Interestingly, PITRM1 deficient mice show an age-dependent accumulation of amyloid precursor protein (APP) and Aβ deposits within the brain ^2^, suggesting a link between defects of mitochondrial proteostasis and adult-onset neurodegenerative dementia. However, due to the embryonic lethality observed in complete PITRM1 knockout mice, the exact role of PITRM1 in brain homeostasis and disease could not be studied ^2^. Since PITRM1 is involved in the degradation of MTS, as well as Aβ peptides ^8^, the pathomechanisms of neurodegeneration linked to loss of PITRM1 function could in principle be due to either accumulation of Aβ peptides in mitochondria, MTS-driven toxicity, or a combination of both. Identifying the mechanisms that lead to neurodegeneration in primary mitochondrial diseases characterized by defects of mitochondrial proteostasis, such as PITRM1-linked neurological syndrome, may help elucidate the long-debated, still unresolved, involvement of altered mitochondrial function in neurodegenerative dementia. To examine PITRM1-related pathogenetic mechanisms, we generated PITRM1-knockout human induced pluripotent stem cells (iPSCs) and examined the role of mitochondrial function and proteostasis using 2D neuronal and 3D brain-organoid model systems.

## Methods

### Generation of PITRM1 knockout human iPSCs

Control iPSCs (from a female non affected control, 80 years) used in this study were previously generated and characterized ^12^. All cells used in the study were derived from patients who signed an informed consent approved by The Ethics Committee of the Medical Faculty and the University Hospital Tübingen. iPSCs were kept in culture in hESC medium ^12^. SgRNAs targeting exon 3 and 4 of PITRM1 gene were designed using CRISPR Guide Design Tools (former www.crispr.mit.edu) and purchased from Metabion International AG (exon 3 top AGGAGCCAGGTATTTACACC, exon 3 bottom GGTGTAAATACCTGGCTCCT, exon 4 top TTGAGCATACCGTCCTTTGT, exon 4 bottom ACAAAGGACGGTATGCTCAA). SgRNAs were cloned into the pSpCas9(BB)-2A-Puro plasmid containing the sgRNA scaffold and puromycine-resistance under the U6 promoter (Addgene plasmid #48139). Colonies that successfully integrated sgRNA into backbone plasmid were screened and confirmed by Sanger sequencing using U6 promoter region primer. SgRNA/Cas9 plasmid was delivered into cells using Nucleofector Amaxa system (Lonza Biosciences). In brief, iPS cells were dissociated with Accutase (Sigma-Aldrich) and 10^6 iPSCs were nucleofected with 6µg of each sgRNA plasmid. Cells were then replated on MEF cells in hESC medium, without P/S, supplemented with 10µM ROCK Inhibitor Y-27632 2HCl. After recovery, cells were replated at density 500 cells/cm^2 for single cell subcloning. After recovery, iPSCs were clonally expanded and the genomic deletion was assessed by PCR and Sanger sequencing (exon 3 Fw TTCAGGCAGAAAAGCCAGTT, exon 4 Rv ACTGAATTCCAGTGGGTGTGC). The screening of possible off-target effects was performed using CRISPR Design Tools. Sequencing primers for off-target effects: (5‘-3‘): PPIL2 NM_148175 Fw CCTCATGCCCTGCTTGACTC, Rv CAGGGAGCACTGTCCCAATTT; NR1D1 NM_021724 Fw CAAACGAGCACACACCACAG, Rv GCTGCCCCCTTGTACAGAAT; KDEL2 NM_001100603 Fw TTGGTGGTGGTTATGCCTCA, Rv ACCACCAGAAACTCCACTCG; FAM120A NM_014612 Fw TCCTGCGGTTCTTGTCCTCTA, Rv GCATGAATGTGTCTTCTCTGGC; TTLL2 NM_031949 Fw GTGGGAGGCTGTGTGGTATT, Rv TCAAGTCCCTACCTGTGCCA; SUCO NM_014283 Fw AATCTGGTACTATTCCGATAGCCAA, Rv CCATTCAAACAGGACACTGCTG.

### Cortical neuronal differentiation

For the induction of cortical neurons, we used an embryoid body (EB)-based differentiation protocol, with minor modifications ^13^. iPSC colonies were manually picked and grown for 4□d in EB media (20% KO serum replacement in DMEM/F12 medium, 1% NEAA, 1% P/S). On day 5, EBs were plated onto Matrigel-coated (Corning) plate and grown for 4 more□days in N2B27 media (DMEM/F12, 1X N2, 1X B27-RA, 1% NEAA, 1% P/S, 20□ng/mL bFGF). For the first 8□differentiation days, the cells were grown in the presence of 10□μM SB431542 (Ascent Scientific) and 2.5□μM dorsomorphin (Sigma Aldrich). After 8□d, neural rosettes were lifted with Dispase and replated onto matrigel-coated plate and grown in N2B27 media. Secondary or tertiary rosettes were manually dissected to purify neural progenitor cells (NPCs). For cortical differentiation, NPCs were dissociated with Accutase and seeded at a density of 1,000 cells per mm2 on Matrigel-coated plates in neuronal differentiation medium consisting of DMEM/F12, 1X Glutamax, 1X N2, 1X B27-RA, 20□ng/mL BDNF (Peprotech), 20□ng/mL GDNF, 1□mM dibutyryl-cyclic AMP (Sigma), 200□nM ascorbic acid (Sigma). Medium was replaced every other day.

### Cerebral organoids culture and immunohistochemistry

Cerebral organoids were generated and maintained using the protocol described by Lancaster et al. ^14^. Where indicated, cerebral organoids were treated with 500nM ISRIB or 500μM NMN (both from Sigma Aldrich) daily, from DIV 20 to DIV 50 or from DIV45 to DIV50 respectively. For immunostaining, organoids were washed with PBS, fixed in 4% PFA for 15 min and then equilibrated in 30% sucrose in PBS overnight at 4°C. Next day, organoids were embedded in blocks with a mixture of 10% sucrose/7.5% gelatin, snap frozen, and kept at −80°C until cryosections were prepared using Leica CM 1900 instrument with 20 μm thickness. Sections were then permeabilized and blocked with 10% normal goat serum in PBS in 0.5% Triton X-100. Primary antibody incubations were performed at 4°C overnight, followed by three 10 min washes in PBS, and staining with AlexaFluor secondary antibodies (Invitrogen, 1:1000) at room temperature for 1h. Primary antibodies included rat anti-CTIP2 (1:500, Abcam ab18465), rabbit anti-TBR1 (1:500, Abcam ab31940), mouse anti-β-III-tubulin (1:1000, Sigma-Aldrich T8328), rabbit anti-β-III-tubulin (1:1000, Biolegend, Previously Covance # PRB-435P), chicken anti-MAP2 (1:3000, Biolegend PCK-554P), mouse anti phospho-tau (AT8 1:500, Thermofisher #MN1020), mouse anti-APP (1:500, Santa Cruz Biotechnology sc53822), mouse anti-Ubiquitin (1:100, Merck MAB1510), mouse anti-Caspase 3 (1:500, Cell Signaling Technology 9664T). For thioflavin T immunostaining, sections were stained with 10μM thioflavin T (Sigma-Aldrich T3516) for 15□min at room temperature and washed 3 times in PBS. Cell nuclei were stained with DAPI and slides were mounted with DAKO mounting medium (Agilent S302380-2). Images were acquired using Leica TCS SP8 confocal microscope (Leica Microsystems). For image analysis, mean of fluorescence intensity per image was calculated with Image J. For each condition, 10-15 images were acquired from at least 5 organoids from three independent experiments (cultures).

### Measurement of mitochondrial membrane potential

For measurement of mitochondrial membrane potential, neurons were split at a concentration of 2×10^5 cells/well on 96-wells pre-coated with Matrigel. Cells were washed once with HBSS (Invitrogen) following incubation with 200 nM Tetramethylrhodamine Methylester Perchlorat (TMRM) (Invitrogen) in HBSS with 1% BSA for 30 min at 37°C. Cells were then washed twice with HBSS, detached using Accumax (Invitrogen) and kept in HBSS with 1% BSA (Sigma) for FACS analysis using MaxQuant Analyzer 10 (Miltenyi Biotec). Analysis was performed using the MACSQuantify 2.6 software (Miltenyi Biotec).

### Measurement of mitochondrial Reactive Oxygen Species (mtROS)

For measurement of mtROS production of iPSC-derived neurons by flow cytometry, cells were pre-incubated with N2 medium for 48 hrs. Then, 1×10^6 cells were washed once with HBSS, incubated with 5µM of the superoxide indicator MitoSOX Red (Invitrogen) for 30 min at 37°C, washed twice with HBSS, dissociated using Trypsin (Invitrogen) and resuspended in 200μl of HBSS and 1% BSA. Cytofluorimetric analysis was performed using MACSQuant Analyzer 10 (Miltenyi Biotec).

### Seahorse XF^e^96 Metabolic Flux Analysis

Oxygen consumption rate (OCR) was analyzed using an XFp Extracellular Flux Analyzer (Agilent). iPSC-derived neurons were plated on XFp microplates (Agilent) at a density of 70,000 per well and grown in N2 medium for 48 hrs before the experiment. Measurement of neuronal oxidative consumption rate was performed in freshly prepared medium consisting of phenol-free DMEM,1 mM Natrium Pyruvate, 2mM Glutamine and 10mM glucose with pH adjusted to 7,4. Mitochondrial function was evaluated after subsequent injection of 10μM oligomycin, 10 μM carbonyl cyanide p-trifluoromethoxyphenylhydrazone (CCCP) and 2µM Antimicyn A / 1µM Rotenone (all Sigma-Aldrich). For each condition 3 measurements, lasting 5 minutes each, were performed: after each injection, OCR was measured for 2 minutes, the medium was mixed for 2 min, let rest for 1 min, and then respiration was measured again. After measurement, values were normalized to cell number by counting DAPI stained nuclei using a high-content cell analyzer (BD Bioscience, Pathway 855).

### Quantitative RT-PCR

mRNA was isolated using an RNA isolation kit (Qiagen). Following the reverse transcription reaction using the QuantiTect Reverse Transcription kit, quantitative PCR reaction was performed using SYBR GREEN (all Qiagen) and monitored with a Viia7 Real time PCR system (Applied Biosystems). The expression level of each gene was normalized to the housekeeping gene ribosomal protein large P0 (Rplp0). Fold-changes in gene expression were calculated using the 2^−DDCT^ method, based on biological reference samples and housekeeping genes for normalization.

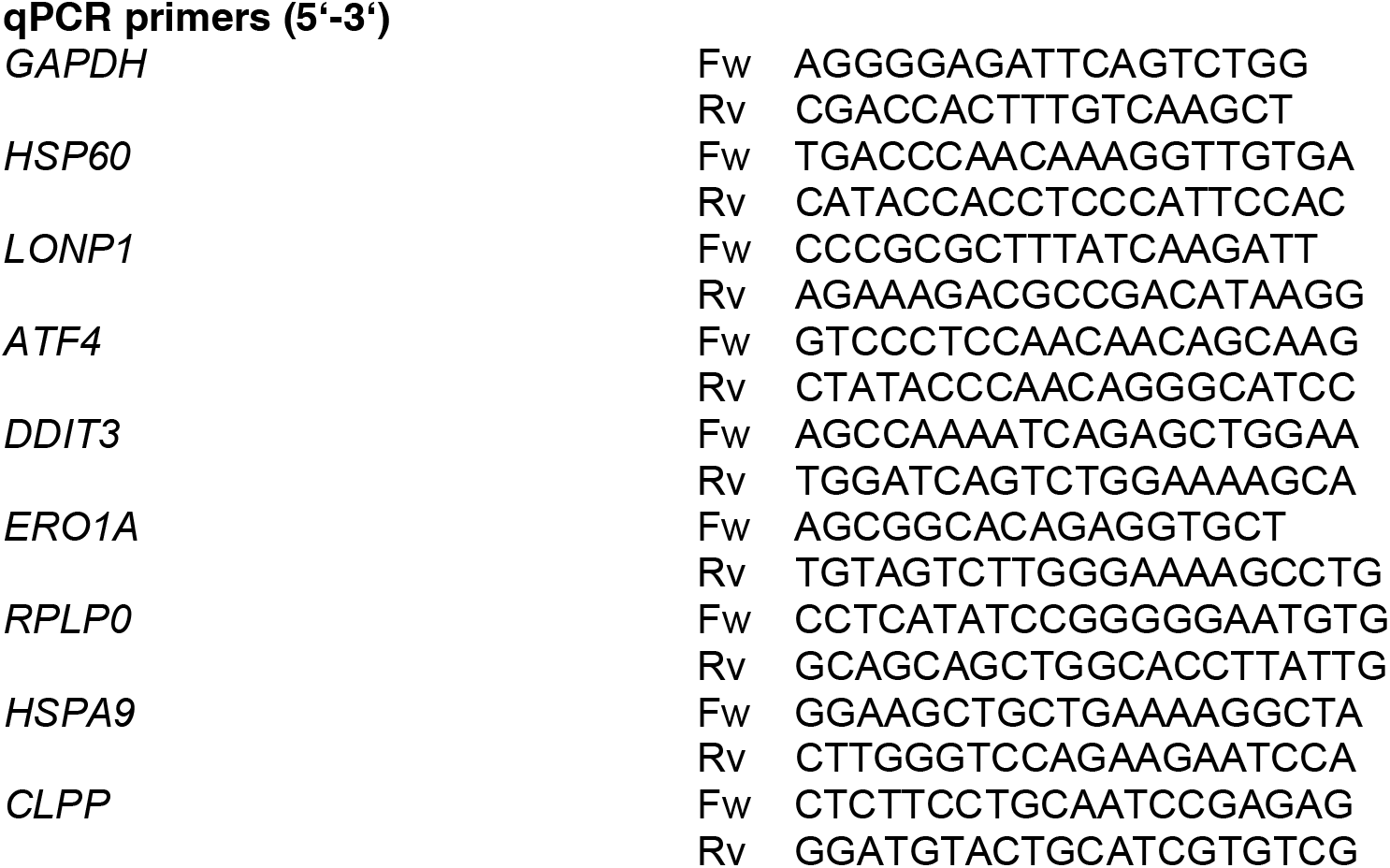
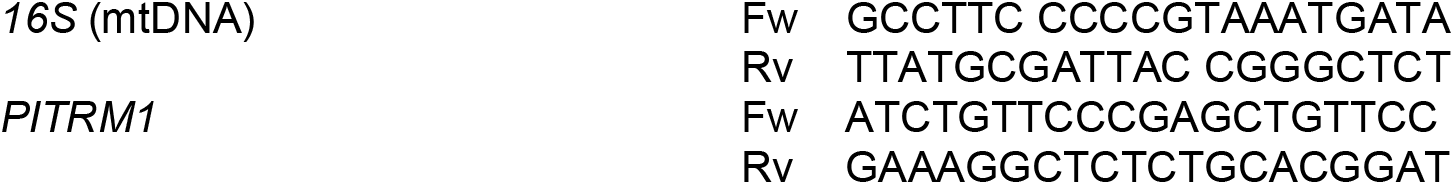

### Mitochondrial isolation

The mitochondrial isolation was performed using Qproteome® Mitochondria Isolation Kit (Qiagen) according to manufacturer’s protocol.

### Western blot

Proteins were extracted using Tris-buffered Saline (TBS) with 0.5% NP40 protein extraction buffer containing protease and phosphatase inhibitors (Roche), on ice, following centrifugation at 14.000 rpm and 4°C for 15min. The protein concentration of the supernatant was determined by BCA (Pierce). In total, 15–30□μg of the protein lysate was loaded on polyacrylamide gel (density ranged from 7.5% to 15%, depending on respective protein molecular weight) and transferred on a PVDF membrane (Millipore). Blots were blocked with 5% milk powder or 5% BSA in TBS + 0.1% Tween-20 (TBST) and incubated with primary antibodies in milk or BSA blocking solution overnight at 4□°C. This step was followed by incubation with corresponding HRP-conjugated secondary antibodies (Sigma Aldrich) for 1 h at room temperature. Visualization of proteins was done by using Amersham ECL Western Blotting Detection Reagent and Amersham Hyperfilm (both GE Healthcare). Densitometric analysis of proteins was performed by ImageJ software. Primary antibodies included: rabbit anti-LC3B (1:500, Cell Signaling Technologies #2775); mouse anti-APP 6E10 (Abeta 1-16) (1:1000, Biolegend #803004); rabbit anti-PITRM1 (1.1000; Atlas Antibodies #HPA006753); rabbit anti-Frataxin (1:1000, Abcam #ab175402); mouse anti-Total OXPHOS human cocktail (1:1000, Abcam #ab110411); mouse anti-ubiquitin (1:5000, Millipore #MAB1510); mouse anti-HSPA9 (1:3000, Santa Cruz Biotechnologies #sc-133137); mouse anti-HSP60 (1:3000, Santa Cruz Biotechnologies #sc-271215); rabbit anti-LONP1 (1:3000, Proteintech #15440-1-AP); mouse anti-tau (1:1000, HT7, Thermofisher #MN1000); mouse anti phospho-tau PHF-6 (Thr231) (1:1000, Thermofisher #35-5200); mouse anti-β-Actin (1:20000, Sigma Aldrich #A5441); mouse anti-VDAC1 (1:1000, Santa Cruz Biotechnologies #sc-390996).

### Autophagy studies

Where indicated, cells were treated with NH_4_Cl (20□mM) and leupeptin (200□μM) (EMD, Millipore) for 4□h. LC3-II and LC-I levels were quantified by densitometry and normalized to β-actin. LC3 flux was quantified by dividing the levels of LC3-II after treatment with lysosomal inhibitor for 4h, by the levels of LC3-II without treatment.

### Immunofluorescence

Cells were fixed in 4% paraformaldehyde (PFA) in PBS (w/v) for 10 min, rinsed with PBS and blocked by 10% normal goat or donkey serum (NGS/NDS) in PBST (PBS + 0.1% TritonX-100) for 60 min. Cells were then incubated with primary antibodies in 10% NGS/NDS in PBST over night at 4°C following 1h incubation at 24°C with appropriate Alexa488/568 coupled secondary antibodies (1:1000, Invitrogen). Cell nuclei were stained with DAPI and final images were acquired using Leica TCS SP8 confocal microscope (Leica Microsystems) and analyzed using Fiji software. Primary antibodies included: rabbit anti-LC3B (1:500, Cell Signaling Technologies #2775); mouse anti-CTIP2 (1:500, Abcam ab18465), rabbit anti-TBR1 (1:500, Abcam ab31940), mouse anti-β-III-tubulin (1:1000, Sigma-Aldrich T8328-25UL), and anti-Caspase 3 (1:500, Cell Signaling Technology 9664T). LC3 particle number in β-III-tubulin positive cells was quantified with the “analyse particles” plug-in in ImageJ (NIH). Caspase 3 was quantified with mean fluorescence intensity in Image J (NIH). Quantification was carried out on, at least, 50 cells per condition, from three independent experiments.

### LDH assay

The LDH assay (Promega) was performed as per the manufacturer’s instructions.

### Amyloid-beta species measurement

For amyloid-beta species measurement in iPSC-cortical neurons, 10^6 cells were plated into a one well of 12-well plate and cell supernatant was collected after 5 days, snap frozen and stored at −80 °C until analysis. Cell pellet was collected and lysed for protein concentration determination and used for value normalization. Conditioned medium from at least 3 technical replicates was collected in each experiment. For amyloid-beta species measurement in cerebral organoids, individual organoids were re-plated into low attachment 96-well plates on an orbital shaker. Supernatant was collected after 5 days, snap frozen in liquid nitrogen and stored at −80 °C until analysis. After collection of supernatant, individual organoids were lysed and protein concentration measured with BCA for value normalization. The concentrations of Aβ40 (Aβx–40) and Aβ42 (Aβx–42) in the samples were measured on a Sector Imager 6000 using an electrochemiluminescence-based immunoassay, V-PLEX Ab Peptide Panel 1 (6E10) Kits (Meso Scale Discovery, Gaithersburg, MD, USA) according to manufacturer’s instructions. All samples were thawed on ice and diluted 1:2 in buffer (Diluent 35, Meso Scale Discovery) before incubation. Neural differentiation medium without B27/N2 supplements was used as negative control for each measurement. Human CSF samples were used as internal references on each plate. At least 8 organoids were used for each measurement. Every sample was tested in duplicate (and excluded if the coefficient of variance (CV) was above 20%). Data analysis was run with the MSD DISCOVERY WORKBENCH software version 2.0.

### Elisa

Individual cerebral organoids were homogenized in ice-cold RIPA buffer containing protease and phosphatase inhibitors (Roche), following centrifugation at 14.000 rpm and 4°C for 10min. The protein concentration of the supernatant was determined by BCA (Pierce). Total and phospho-tau levels were assessed, in equal protein amounts, using ELISA assays (both Invitrogen, KHB0041 and KHO0631, respectively) according to manufacturer’s instructions.

### Statistical analysis

The Statistical Package GraphPad Prism version 7 and 8.3 (GraphPad Software) was used to analyze the data. Statistical significance was evaluated using two-tailed Student’s t-test. Data are expressed as mean + S.E.M. or S.D. as indicated.

## Results

### Mitochondrial dysfunction in PITRM1 knockout iPSC-derived neurons

In order to overcome the limitation of the embryonic lethality previously observed in PITRM1 knockout mice ^2^ and examine the mechanistic link between PITRM1 deficiency and neurodegeneration in a model that more closely resembles human disease, we generated PITRM1 knockout (PITRM1^−/−^) human iPSCs using CRISPR/Cas9 endonuclease-mediated gene editing. Appropriate sgRNAs targeting exon 3 and exon 4 were designed to introduce a frameshift deletion resulting in the complete knockout of PITRM1 protein (Supplementary Fig. 1A). Several homozygous clones were generated, and two fully characterized clones were selected and used for further analysis (Supplementary Fig. 1B-D). To address the impact of PITRM1 on the function of human neurons, we differentiated PITRM1^+/+^ and PITRM1^−/−^ iPSCs into neuronal cultures that were enriched for cortical neurons and we assessed neuronal cultures at 35 and 65 days *in vitro*. Both PITRM1^+/+^ and PITRM1^−/−^ iPSCs efficiently generated cortical neurons (Fig. 1A). Western blot analysis confirmed the complete absence of PITRM1 protein in PITRM1^−/−^ iPSC-derived neurons (Fig. 1B). No overt cell death was observed in neuronal cultures, as assessed by LDH assay at 35 and 65 days *in vitro* (Supplementary Fig. 1E) and cleaved Caspase 3 staining (data not shown). PITRM1 deficiency leads to the accumulation of non-degraded MTS sequences that in turn impair the processing of presequence proteins by the peptidase MPP ^8^. Since MPP is also involved in the maturation of the human frataxin precursor ^15^, we examined frataxin maturation by immunoblotting. A decreased ratio of processed, mature to immature frataxin was detected in PITRM1^−/−^ iPSC-derived neural precursor cells (NPCs) and neurons (Supplementary Fig. 1F, G), indicating the impaired function of MPP and defects of mitochondrial of mitochondrial presequence processing. Since MTS peptides can bind to the membrane and perturb the mitochondrial electrochemical gradient, we evaluated the effect of PITRM1 deficiency on mitochondrial membrane potential and respiratory oxidative activity. Mitochondrial membrane potential was significantly reduced in PITRM1^−/−^ neurons compared to isogenic PITRM1^+/+^ neurons (Fig. 1C). However, mitochondrial reactive oxygen species (mtROS) were not significantly altered in PITRM1^−/−^ neurons (Fig. 1D). Similarly, no significant difference in the respiratory oxidative activity was observed between PITRM1^+/+^ and PITRM1^−/−^ neurons (Fig. 1E). Western blot analysis revealed a significant increase in the level of Complex II respiratory chain complex subunits in PITRM1^−/−^ neurons (Fig. 1F, G). However, no significant difference for the levels of all other respiratory chain complex subunits was detected in PITRM1^−/−^ neurons (Fig. 1F, G).

**Figure 1.**
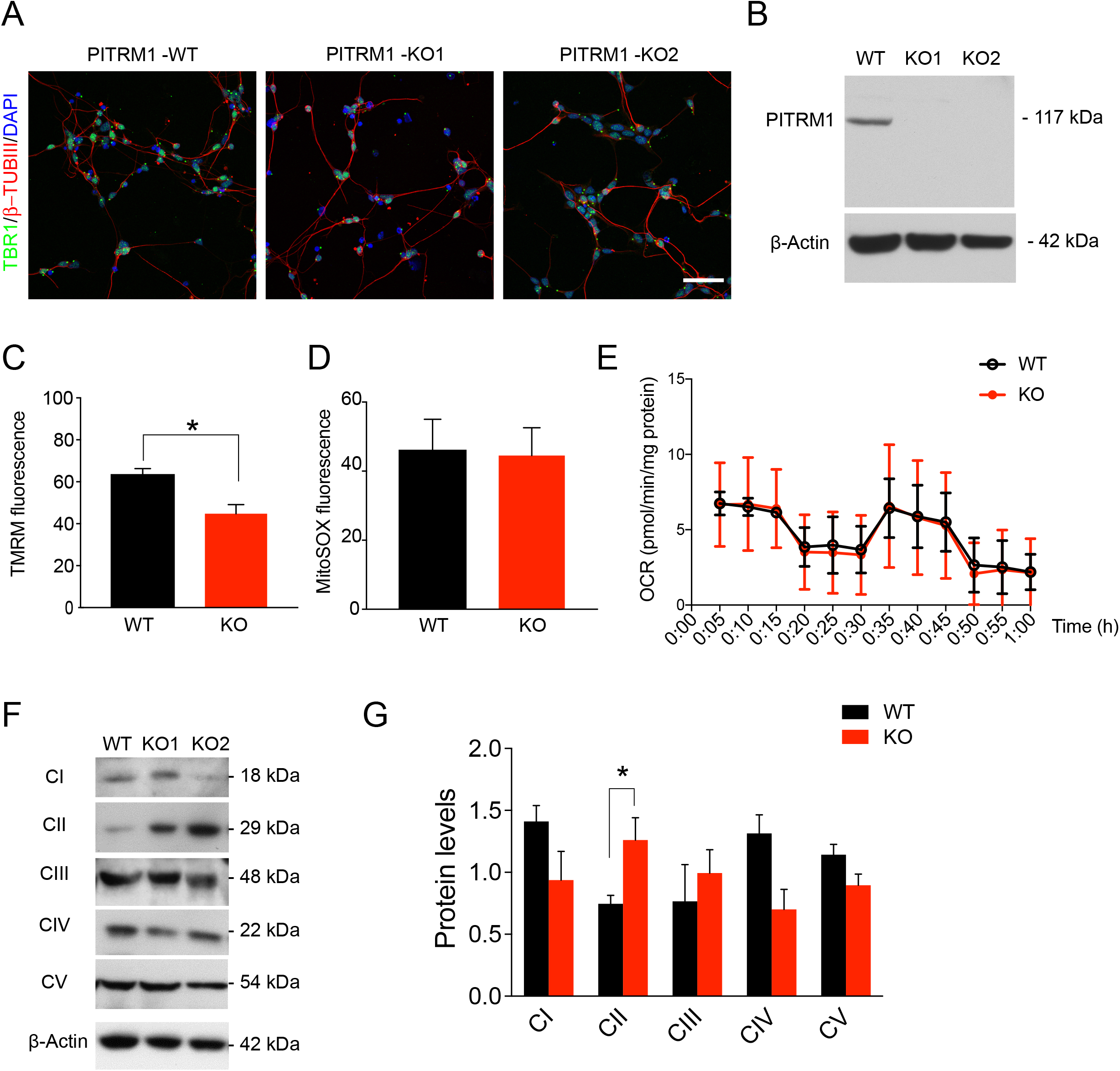
Mitochondrial dysfunction in PITRM1^−/−^ iPSC-derived cortical neurons. Control PITRM1^+/+^ (WT) and isogenic PITRM1^−/−^ (KO) iPSCs were differentiated into cortical neurons. **(A)** Immunostaining of indicated differentiated iPSC cultures at day *in vitro* 35. Cells were stained for TBR1 (green) and β-III Tubulin (β-TUBIII, red). Nuclei were counterstained with DAPI (blue). Scale bars, 50 μm. **(B)** Representative western blot for PITRM1, showing absence of PITRM1 in human PITRM1^−/−^ iPSC-derived cortical neurons. **(C)** Mitochondrial membrane potential in isogenic PITRM1^+/+^ and PITRM1^−/−^ iPSC-derived neurons, as determined by a tetramethylrhodamine methyl ester (TMRM) assay and flow cytometry analysis. Data are expressed as percentage of TMRM positive cells (mean + SEM; * p<0.05, two-tailed t test, n=4). **(D)** Mitochondrial reactive oxidative species level as analyzed by MitoSOX labeling in isogenic PITRM1^+/+^ and PITRM1^−/−^ iPSC-derived neurons (mean + SEM, n=5). **(E)** Oxygen consumption rate (OCR) of PITRM1^+/+^ and PITRM1^−/−^ iPSC-derived neurons. Data are normalized to protein content (mean ± SD, n=6). **(F, G)** Western blot analysis of OXPHOS complex protein levels in PITRM1^+/+^ and PITRM1^−/−^ iPSC-derived neurons. Representative blot is shown in (F) and the quantification is shown in (G) (mean + SEM; * p<0.05, two-tailed t test, n=4).

### PITRM1^−/−^ iPSC-derived cortical neurons show induction of mitochondrial stress response and enhanced mitophagy

Mitochondrial stress response has been identified as a common signature in several neurodegenerative as well primary mitochondrial diseases ^16-19^. Thus, we examined the expression levels of genes involved in mitochondrial unfolded protein response (UPR^mt^) and, more generally, in the mitochondrial integrated stress response pathway (mtISR). PITRM1^−/−^ iPSC-derived cortical neurons exhibited a significant induction of UPR^mt^/mtISR transcripts (*ATF4, DDIT3, HSP60, HSPA9, ERO1*) (Fig. 2A). Moreover, gene expression of the mitochondrial proteases, *LONP1 and CLPP*, was significantly upregulated in PITRM1^−/−^ neurons compared to controls (Fig. 2A). In line with gene expression data, we found an increase in protein expression of the chaperones HSPA9 and HSP60, and the mitochondrial protease LONP1 (Fig. 2B). These data indicate that the accumulation of MTS, due to PITRM1 deficiency, leads to a strong upregulation of the ISR pathway. Since, the ISR has been shown to activate autophagy ^20, 21^, we assessed the autophagosome content by immunostaining for endogenous light chain type 3 protein (LC3), a marker of autophagosomes. Our analysis revealed a significant decrease of LC3-positive vesicles in PITRM1^−/−^ neurons compared to isogenic PITRM1^+/+^ neurons (Fig. 2C, D). These results were confirmed by Western blot, showing decreased basal levels of LC3-II in PITRM1^−/−^ neurons (Fig. 2E). However, inhibition of lysosomal degradation by leupeptin and ammonium chloride revealed that the autophagic flux was significantly increased in PITRM1^−/−^ neurons, thus confirming autophagy activation (Fig. 2E, F). To assess whether the observed increase in autophagic flux leads to an enhanced turnover rate of mitochondria by autophagy, we purified mitochondria from PITRM1^+/+^ and PITRM1^−/−^ iPSC-derived cortical neurons. Immunoblotting of purified mitochondria showed increased ubiquitination of PITRM1^−/−^ neurons compared to PITRM1^+/+^ neurons (Fig. 2G, H), suggesting their targeting to lysosomes and increased mitochondrial clearance ^22^. In line with these results, mitochondrial content was lower in PITRM1^−/−^ neurons, as shown by a reduced mitochondrial to nuclear DNA ratio (Fig. 2I). Taken together, these results suggest that loss of PITRM1 function enhances mitochondrial clearance.

**Figure 2.**
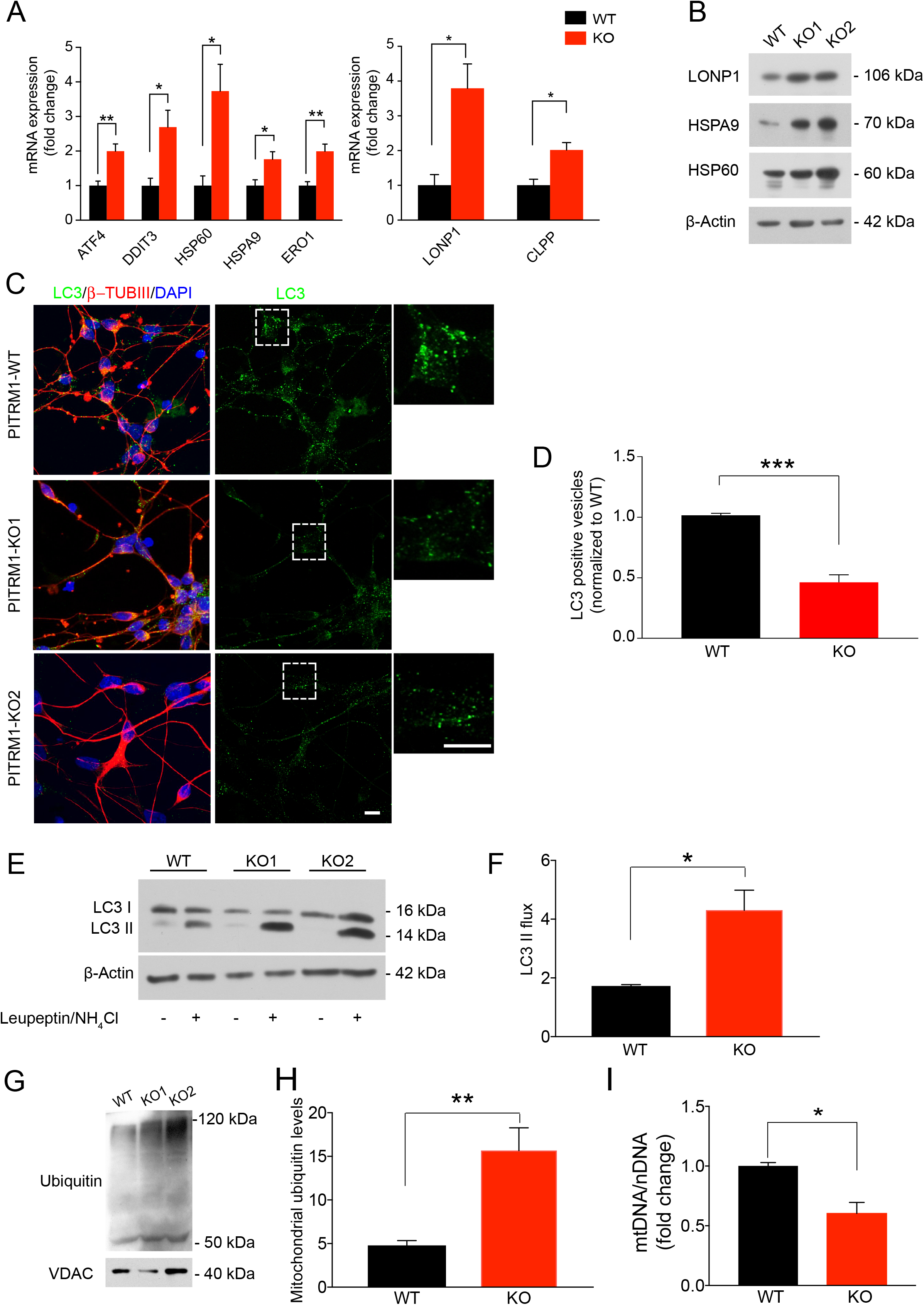
PITRM1^−/−^ iPSC-derived neurons show induction of UPR^mt^ and enhanced mitophagy. **(A)** Gene expression levels of mitochondrial stress response genes in PITRM1^+/+^ and PITRM1^−/−^ iPSC-derived cortical neurons (mean + SEM; ** p<0.01, * p<0.05, two-tailed t test, n=5). **(B)** Representative Western blots of the mitochondrial chaperones HSPA9, HSP60 and the mitochondrial protease LONP1 in PITRM1^+/+^ and PITRM1^−/−^ iPSC-derived cortical neurons. **(C)** Immunostaining of PITRM1^+/+^ and PITRM1^−/−^ iPSC-derived cortical neurons for LC3 (green) and β-TUBIII (red). Nuclei were counterstained with DAPI (blue). Scale bars, 10 μm. **(D)** Number of LC3-positive vesicles per β-TUBIII positive cell relative to control neurons (mean + SEM; *** p<0.001, two-tailed t test, n=3). **(E)** Western blot analysis for LC3 in PITRM1^+/+^ and PITRM1^−/−^ iPSC-derived neuronal cultures, untreated (−) or treated with 200□μM leupeptin and 20□mM NH_4_Cl for 4□h (+). **(F)** Quantification of LC3 flux normalized to WT (mean + SEM; * p<0.05, two-tailed t test, n=4). **(G)** Representative Western blot of isolated mitochondria from PITRM1^+/+^ and PITRM1^−/−^ iPSC-derived neurons with an antibody for ubiquitination and VDAC as loading control. **(H)** Quantification of mitochondrial protein ubiquitination levels in PITRM1^+/+^ and PITRM1^−/−^ iPSC-derived neurons (mean + SEM; ** p<0.01, two-tailed t test, n=3). **(I)** mtDNA content was measured as mitochondrial (*16S*) to nuclear (*RPLP0*) DNA ratio by qRT-PCR (mean + SEM; * p<0.05, two-tailed t test, n=3).

### PITRM1^−/−^ iPSC-derived neurons show accumulation of APP and increase in extracellular Aβ peptides levels

To examine the impact of PITRM1 activity on Aβ pathology, we first assessed the levels of APP by Western blot. APP protein levels were found significantly increased in PITRM1^−/−^ neurons compared to PITRM1^+/+^ neurons (Fig. 3A). Next, supernatant from PITRM1^+/+^ and PITRM1^−/−^ cortical neurons was analyzed using the Meso Scale Discovery immunoassay for human Aβ peptides. PITRM1^−/−^ neurons showed significantly higher levels of Aβ40 and Aβ42 peptides as well as an increased Aβ42/Aβ40 ratio compared to control samples (Fig. 3B). Remarkably, Aβ peptides were not detected in mitochondrial extracts from PITRM1 deficient neurons. Next, we explored the mechanisms that link loss of PITRM1 function with alteration of APP metabolism. Since UPR^mt^ may have an impact on ubiquitin-dependent protein turnover ^23^, we examined the levels of ubiquitinated proteins by Western blot. We observed that PITRM1^−/−^ display increased levels of ubiquitinated proteins (Fig. 3C, D), suggesting defects in cellular proteostasis.

**Figure 3.**
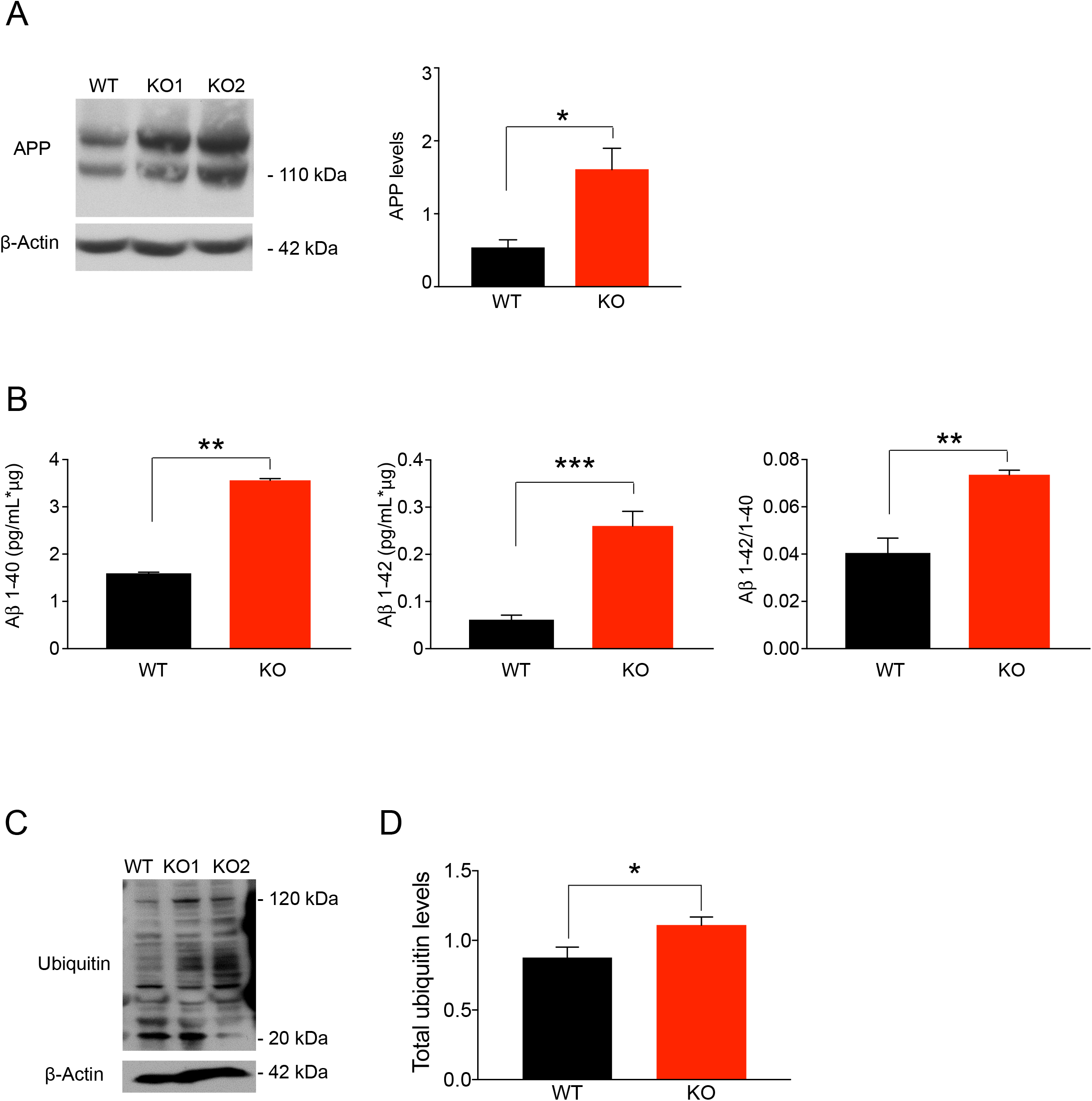
PITRM1^−/−^ iPSC-derived neurons show accumulation of APP and increased levels of Aβ peptides. **(A)** Representative Western blots of APP in PITRM1^+/+^ and PITRM1^−/−^ iPSC-derived cortical neurons. Quantification of APP levels normalized to loading control (mean + SEM; * p<0.05, two-tailed t test, n=5). **(B)** Quantification of Aβ species and Aβ42/Aβ40 ratio in the supernatant of PITRM1^+/+^ and PITRM1^−/−^ iPSC-derived cortical neurons at day in vitro 35, as performed by Meso Scale immunoassay (mean + SEM; ** p<0.01, *** p<0.001, two-tailed t test, n=4). **(C)** Representative Western blot of PITRM1^+/+^ and PITRM1^−/−^ iPSC-derived neurons of total ubiquitinated proteins levels. **(D)** Quantification of protein ubiquitination level in PITRM1^+/+^ and PITRM1^−/−^ iPSC-derived neurons (mean + SEM; * p<0.05, two-tailed t test, n=4).

### PITRM1^−/−^ cerebral organoids exhibit main features of AD pathology and induction of mitochondrial stress response

Despite APP accumulation and increased Aβ42/Aβ40 ratio, 2D iPSC-derived neuronal cultures did not show Aβ aggregates nor tau pathology or overt cell death. To further investigate the mechanisms of PITRM1 neurotoxicity in a model that better resembles the human disease, we developed cerebral organoids from PITRM1^+/+^ and PITRM1^−/−^ iPSCs and cultured them over a broad time range (1-6 months). Cerebral organoids derived from PITRM1^+/+^ and PITRM1^−/−^ iPSCs displayed similar sizes and cortical layering characteristics (Fig. 4A and Supplementary Fig. 2A). Next, we examined whether PITRM1^−/−^ cerebral organoids develop AD-like neurodegenerative features. Western blotting reveled increased APP levels and tau hyperphosphorylation in 2-months old PITRM1^−/−^ cerebral organoids (Fig. 4B-E). Furthermore, immunoassay measurements showed a higher Aβ40, Aβ42 and Aβ42/Aβ40 ratio in PITRM1^−/−^ cerebral organoids compared to controls (Supplementary Fig. 2B). Similarly, immunofluorescence staining showed increased APP and phospho-tau levels in PITRM1^−/−^ organoids compared to PITRM1^−/−^ controls starting at 2 months (Fig. 4F, G, I). No further increase of APP levels or tau hyperphosphorylation were observed at later time points (Supplementary Figure 2C). Next, we stained 1-, 2-, and 6-month old PITRM1^+/+^ and PITRM1^−/−^ cerebral organoids, for cleaved caspase-3. The number of cleaved caspase-3 positive cells in the neuroepithelial layers was higher in PITRM1^−/−^ cerebral organoids than controls, starting at 2 months (Fig. 4H, I and Supplementary Fig. 2D), suggesting a higher extent of cell death. No further increase of cell death was observed at later time points (Supplementary Figure 2D). To analyze ubiquitin-dependent protein turnover, an immunostaining against ubiquitinated proteins was performed. PITRM1^−/−^ organoids display increased levels of ubiquitinated proteins (Fig 4J, K). Thioflavin T positive aggregates were detected in cerebral organoids generated from PITRM1^−/−^ iPSCs, indicating that protein deposits are organized into amyloid-like aggregates, similar to the ones observed in AD plaques (Supplementary Fig. 2E). Next, we examined the expression levels of genes involved in UPR^mt^ in 2-month old PITRM1^+/+^ and PITRM1^−/−^ cerebral organoids. PITRM1^−/−^ cerebral organoids exhibited a significant induction of UPR^mt^ transcripts (*ATF4, DDIT3, HSP60, HSPA9, ERO1*) (Fig.4L). Moreover, gene expression of the mitochondrial proteases, *LONP1* and *CLPP*, was significantly upregulated in PITRM1^−/−^ cerebral organoids compared to controls (Fig.4L).

**Figure 4.**
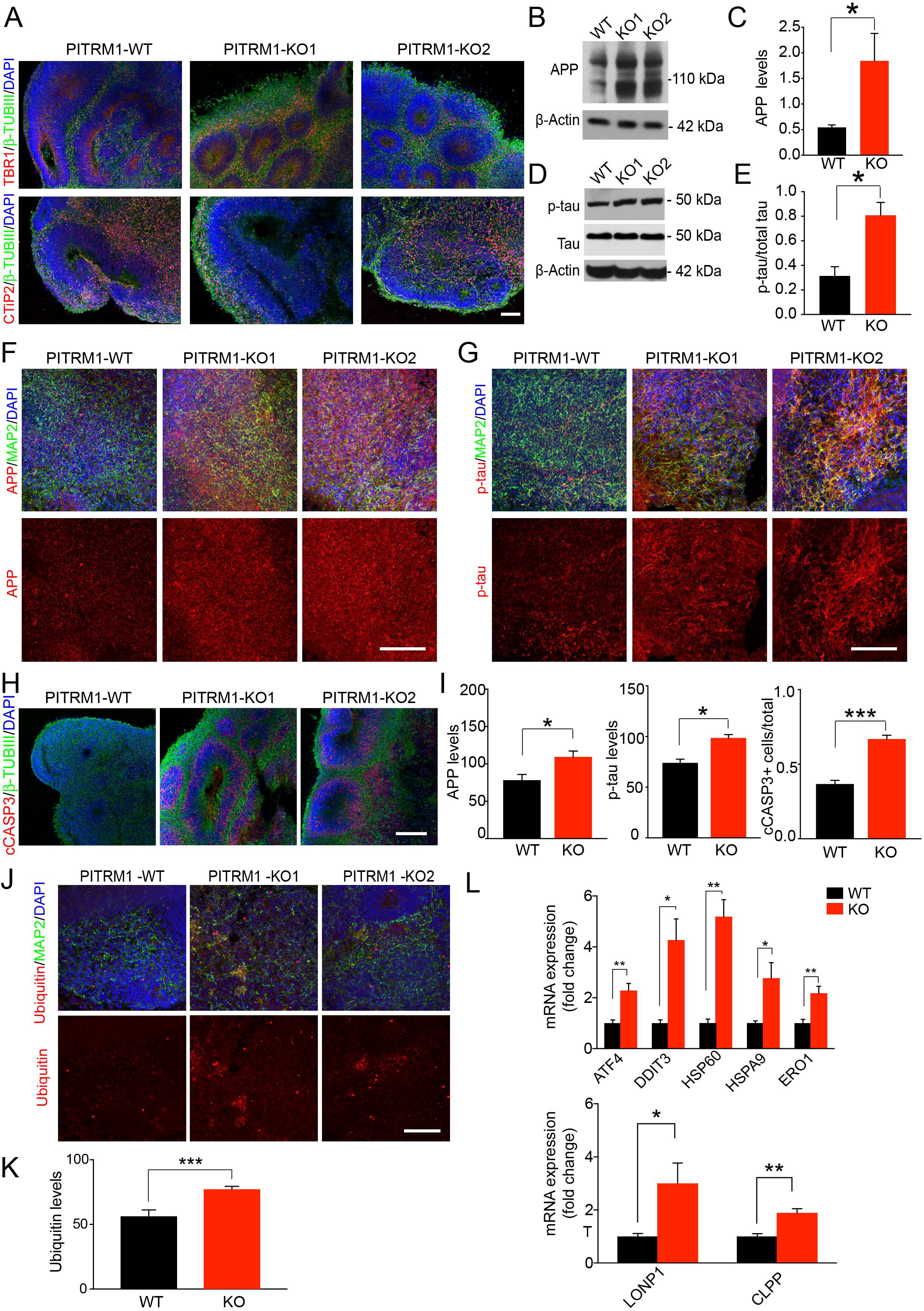
PITRM1^−/−^ cerebral organoids display main pathological features of AD pathology and induction of mitochondrial stress response. **(A)** Generation and characterization of cerebral organoids from PITRM1^+/+^ and PITRM1^−/−^ iPSCs. Immunostaining for β-TUBIII (green), TBR1 (red), and CTIP2 (red) of 2-month old cerebral organoids. Cell nuclei were counterstained with DAPI (blue). Scale bar, 100µm. **(B)** Western blot of APP in 2-month old PITRM1^+/+^ and PITRM1^−/−^ cerebral organoids. **(C)** Quantification of APP protein levels in cerebral organoids (mean + SEM; * p<0.05, two-tailed t test, n=5). **(D)** Representative Western blot of phospho-tau (p-tau) and total tau in PITRM1^+/+^ and PITRM1^−/−^ cerebral organoids; total tau and β-Actin were used as loading controls. **(E)** Quantification of phospho-tau protein levels in cerebral organoids relative to loading control total tau/β-Actin (mean + SEM; * p<0.05, two-tailed t test, n=5). **(F, G)** MAP2 (green), APP (red, left panel), and phospho-tau (red, right panel) immunostaining in cerebral organoids. Representative confocal images are shown. Cell nuclei were counterstained with DAPI (blue). Scale bars, 100 μm. **(H)** Immunostaining for β-TUBIII (green) and cleaved Caspase 3 (cCASP3, red) in PITRM1^+/+^ and PITRM1^−/−^ 2-month old cerebral organoids. Cell nuclei were counterstained with DAPI (blue). Scale bar, 100µm. **(I)** Quantification of APP, phospho-tau fluorescent intensity and analysis of the ratio of cCASP3 positive cells relative to the total number cells, measured by DAPI staining, in 2-month old cerebral organoids (mean + SEM; * p<0.05, *** p<0.001, two-tailed t test, n=3-4). **(J)** MAP2 (green) and Ubiquitin (red) immunostaining in cerebral organoids. Representative confocal images are shown. Cell nuclei were counterstained with DAPI (blue). Scale bars, 100 μm. **(K)** Quantification of Ubiquitin fluorescent intensity (mean + SEM; *** p<0.001, two-tailed t test, n=3). **(L)** Gene expression levels of mitochondrial stress response genes in PITRM1^+/+^ and PITRM1^−/−^ cerebral organoids (mean + SEM; * p<0.05, ** p<0.01, two-tailed t test, n=5).

### Inhibition of UPR^mt^ exacerbates Aβ proteotoxicity

Given that UPR^mt^ can extend lifespan in a variety of organisms ^24-27^, we asked whether the induction of UPR^mt^ observed in PITRM1^−/−^ cerebral organoids could act as a protective mechanism against defects of mitochondrial protein maturation and Aβ proteotoxicity. To this end, cerebral organoids were treated daily with ISRIB, a global ISR inhibitor ^28^. First, we examined APP levels and phospho-tau/tau ratio by immunostaining. Consistent with a protective role of UPR^mt^ in our model, ISRIB-treated cerebral organoids showed higher APP and phospho-tau levels compared to controls (Fig. 5A-C). Remarkably, the effect was similar in both PITRM1^+/+^ and PITRM1^−/−^ organoids (Fig. 5A-C). In parallel, ISRIB treatment significantly increased the Aβ42/Aβ40 ratio (Fig. 5D) in both PITRM1^+/+^ and PITRM1^−/−^ organoids. Interestingly, ISRIB-treated cerebral organoids also showed an increase of mitochondrial DNA, suggesting that inhibition of UPR^mt^ leads to a decrease of mitochondrial clearance (Fig. 5E).

**Figure 5.**
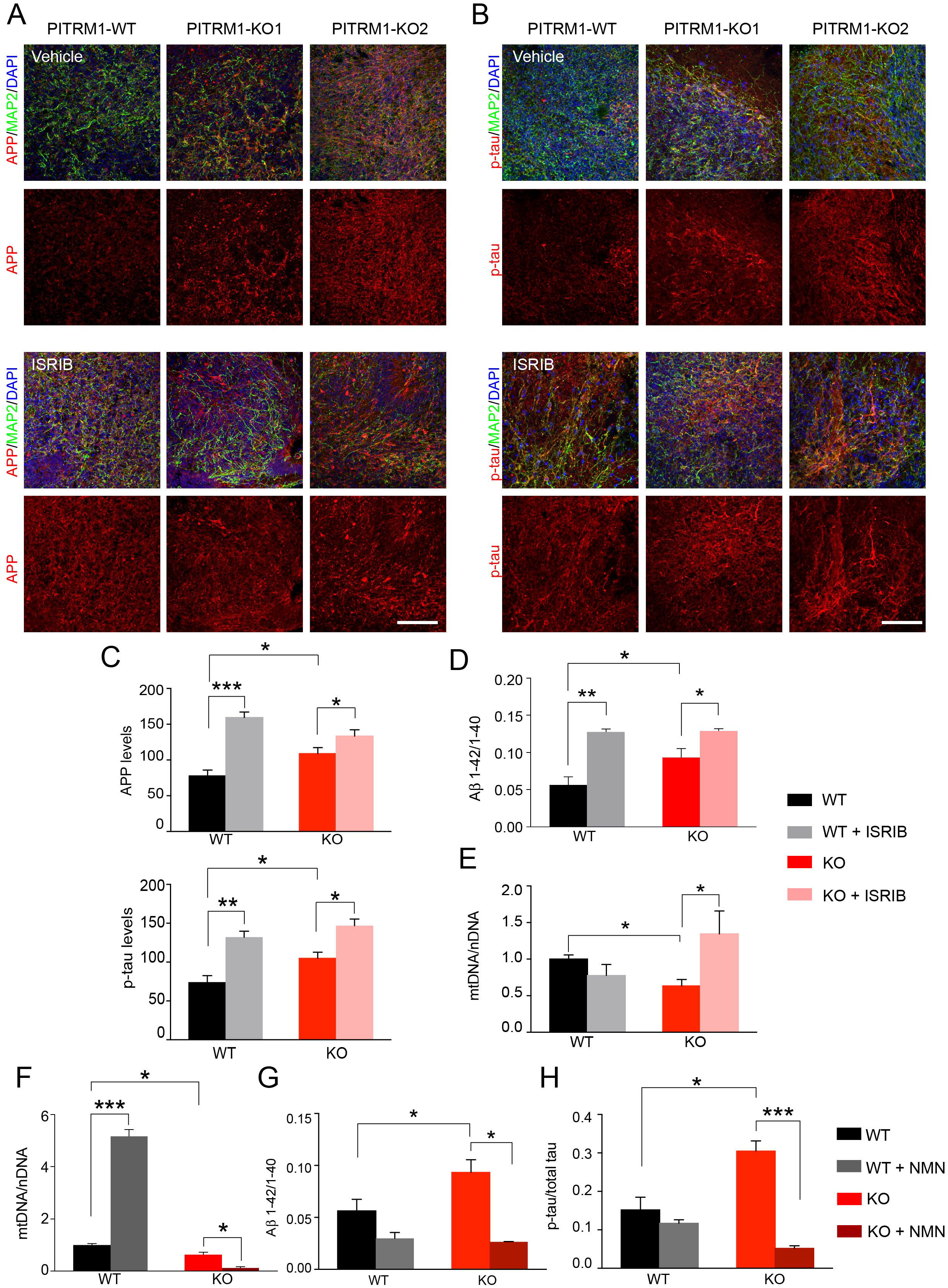
UPR^mt^ and mitophagy exert a protective role in PTRM1^−/−^ cerebral organoids. PITRM1^+/+^ and PITRM1^−/−^ cerebral organoids were treated with 500 nM ISRIB or vehicle. **(A, B)** MAP2 (green), APP (red, in A), and phospho-tau (red, in B) immunostaining in PITRM1^+/+^ and PITRM1^−/−^ cerebral organoids treated with ISRIB or vehicle. Representative confocal images are shown. Cell nuclei were counterstained with DAPI (blue). Scale bars, 100 μm. **(C)** Quantification of APP and phospho-tau fluorescent intensity (mean + SEM; *** p<0.001, ** p<0.01, * p<0.05, two-tailed t test, n=3-4). **(D)** Quantification of Aβ species in supernatant of PITRM1^+/+^ and PITRM1^−/−^ cerebral organoids treated with ISRIB or vehicle, as measured by Meso Scale immunoassay (mean + SD; ** p<0.01, *p<0.05, two-tailed t test, n=5). **(E)** mtDNA content was measured in PITRM1^+/+^ and PITRM1^−/−^ cerebral organoids treated with ISRIB or vehicle as mitochondrial (*16S*) to nuclear (*RPLP0*) DNA ratio by qRT-PCR (mean + SD; * p<0.05, two-tailed t test, n=3). **(F-H)** PITRM1^+/+^ and PITRM1^−/−^ cerebral organoids were treated with 500 μM NMN or vehicle. (F) mtDNA content was measured as mitochondrial (*16S*) to nuclear (*RPLP0*) DNA ratio by qRT-PCR (mean + SEM; *** p<0.001, * p<0.05 two-tailed t test, n=3). (G) Aβ1–42/Aβ1–40 ratio in cerebral organoids measured by Meso Scale immunoassay (mean + SEM; * p<0.05, two-tailed t test, n=3). (H) ELISA assay measuring the levels of total and phospho-tau levels in cerebral organoid homogenates. Protein concentration was measured by BCA and equal amounts of total protein were used (mean + SEM; * p<0.05, *** p<0.001, two-tailed t test, n=3).

### Enhancing mitophagy via NAD+ precursors ameliorates Aβ proteotoxicity

Since defects in mitophagy have been shown to contribute to AD ^29^ and ISRIB-treated cerebral organoids showed decreased mitochondrial clearance and exacerbation of AD-like phenotypes, we investigated whether enhancing mitochondrial clearance ameliorates AD-like phenotype in PITRM1-related mitochondrial neurodegeneration. To this end, PITRM1^+/+^ and PITRM1^−/−^ cerebral organoids were treated with the NAD+ precursor nicotinamide mononucleotide (NMN), which has been shown to ameliorate mitochondrial function and clearance ^30^. While NMN supplementation resulted in increased mtDNA/nDNA ratio in PITRM1^+/+^ organoids, a significant decrease of mitochondrial content was observed in PITRM1^−/−^ organoids after treatment (Fig. 5F). These data suggest a differential role of NAD+ boosters in the physiological and pathological conditions, namely induction of mitochondrial biogenesis in physiological condition and enhancement of mitochondrial clearance in the disease condition. Furthermore, NMN treatment significantly decreased the Aβ42/Aβ40 ratio and phospho-tau/tau levels in PITRM1^−/−^ cerebral organoids as revealed by Meso Scale and Elisa measurements (Fig. 5G, H).

## Discussion

Since the brain is the organ with the highest demand for energy, it comes as no surprise that it also represents the major disease target, both in genetically driven primary mitochondrial diseases, as well as in common age-related neurodegenerative disorders. Despite this evidence, the causal link between mitochondrial demise and neurodegeneration still remains elusive. We have recently reported that pathogenic variants in the nuclear-encoded mitochondrial peptidase PITRM1 result in childhood-onset recessive cerebellar disease leading to a slowly progressive syndrome, characterized by spinocerebellar ataxia, mild intellectual disability, psychiatric manifestations, and cognitive decline ^2, 3^. The clinical picture of these patients is unusual for mitochondrial disease, with a very slow progression of cognitive and psychiatric symptoms from childhood to their late sixties ^2^.

PITRM1^+/−^ mice show a neurological phenotype with presence of Aβ positive plaques in the neuropilum ^2^. However, due to the embryonic lethality of complete PITRM1^−/−2^, the exact mechanisms of PITRM1 in brain function and disease could not be entirely studied. To address these quetions, we have generated a novel human stem cell-based model of loss of PITRM1 function that recapitulates several pathological aspects of human PITRM1-related mitochondrial and adult onset neurodegenerative diseases. Employing iPSC-derived cortical neurons, we found that loss of PITRM1 function leads to a strong induction of mitochondrial stress responses, enhanced autophagic flux and mitochondrial clearance, as well as increased levels of APP, Aβ peptides, and increased Aβ42/40 ratio. Several works have shown the uptake and accumulation of Aβ within the mitochondria in post-mortem brains of AD patients as well as in later stages of disease in APP transgenic mice ^31, 32^. Furthermore, *in vitro* and yeast-based modeling has shown that loss of PITRM1 function results in the incomplete degradation of Aβ in the mitochondria ^2, 8^. Using sub-fractionation methods coupled with a highly sensitive immunoassay, we were unable to detect Aβ peptides in mitochondrial extracts from PITRM1 deficient cortical neurons. We cannot exclude the accumulation of low amount of Aβ peptides within the mitochondria, below the detection limit of this study. However, PITRM1^−/−^ neurons did not show an increase in mitochondrial ROS production or defects in oxidative phosphorylation, which have been shown to be a direct effect of the accumulation of Aβ within mitochondria ^33, 34^. Our data suggests that MTS toxicity and mitochondrial proteostasis imbalance alone can be the first event of the pathogenetic cascade in PITRM1-related neurological syndrome. The mechanisms whereby PITRM1 dysfunction causes APP accumulation may involve proteasome saturation, in response to mitochondrial protein misfolding. The ubiquitin proteasome system (UPS) is also involved in the quality control of mitochondrial proteins, especially the mitochondrial precursor proteins and proteins of the outer membrane ^35^. Supporting an overload of the UPS system, we detected an accumulation of ubiquitinated proteins in PITRM1-deficient neurons and cerebral organoids. Based on these data, we propose that mitochondrial proteotoxic stress, possibly due to the accumulation of non-degraded MTS as a result of PITRM1 dysfunction and accumulation of unprocessed mitochondrial proteins, triggers a cytosolic response with overload and saturation of the proteasome and defects in cytosolic protein degradation.

PITRM1 deficiency led to a strong induction of UPR^mt^ in both 2D and brain organoids model systems. UPR^mt^ is a transcriptional response involving mitochondrial chaperones and proteases activated by mitochondrial dysfunction and defects in mitochondrial protein folding ^36^. The UPR^mt^ is a key cellular quality control mechanism that promotes the maintenance of mitochondrial health and ensures proper cellular functions ^37^. Despite the evidence of UPR^mt^ activation in the ageing and disease brain ^16^, whether and how UPR^mt^ contributes to neurodegenerative processes is unclear. The UPR^mt^ has been proposed to be a double edge sword, with its chronic activation leading to detrimental consequences for cellular and organismal function ^18, 38^. A detrimental role of UPR^mt^ has been demonstrated in animal models of mitochondrial diseases ^18, 38^. However, mitochondrial stress responses have been documented in AD and recent work has shown that enhancing UPR^mt^ provides protections against Aβ proteotoxicity ^17, 39^. In line with a beneficial role of UPR^mt^, PITRM1^−/−^ cerebral organoids treated with ISRIB, an inhibitor of the ISR, showed a significant increase of APP levels, increased Aβ42/Aβ40 ratio, and tau hyperphosphorylation. Based on these data, we propose that PITRM1-related induction of UPR^mt^ is a protective mechanism against proteotoxic stress both at the mitochondrial and cytosolic level. Importantly, our findings indicate that the consequences of chronic mtISR upregulation may vary substantially among different mitochondrial diseases and the underlying molecular defect should be carefully taken into consideration for therapeutic decisions.

Even though UPR^mt^ was activated in both 2D and 3D PITRM1 KO models, only PITRM1 KO cerebral organoids displayed the typical abnormalities observed in the brain of AD patients, including neuronal cell death, tau pathology, and accumulation of protein aggregates, similar to Aβ plaques. On the contrary, despite APP accumulation and increased Aβ42/Aβ40 ratio, we did not detect overt cell death, nor Aβ aggregates or tau pathology in 2D iPSC-derived neuronal cultures. These findings indicate that 3D systems provide a more relevant, compared to 2D, disease model, advantageous in investigating the link between cellular proteostasis and disease. Due to the prolonged culturing conditions, as well as the presence and interaction among different cell types, including glial cells, 3D model systems may promote the development of disease relevant phenotypes, such as protein aggregation and neuronal death ^40^. In respect to the mechanisms, these data also suggest that PITRM1 deficiency triggers compensatory quality control mechanisms both at the cytosolic and mitochondrial level (i.e. induction of UPR^mt^ and autophagy/mitophagy) that ensure the maintenance of cellular proteostasis. However, over time, these mechanisms may be not sufficient to protect neuronal cells against mitochondrial proteotoxicity, as observed in long-term culture cerebral organoids.

Several findings, including the induction of autophagic flux, decreased mtDNA levels, and increase of mitochondrial protein ubiquitination suggest that PITRM1 deficiency leads to increased mitochondrial clearance. It is known that defects in PITRM1 activity lead to impaired MTS processing and accumulation of MTS and precursor proteins that have a toxic effect on mitochondria. In line with this evidence, we report that PITRM1^−/−^ neurons show defects in the maturation of the human frataxin precursor and decreased mitochondrial membrane potential. Enhanced mitophagy could be triggered by mitochondrial depolarization in response to MTS accumulation within mitochondria. Furthermore, UPR^mt^, and in general the ISR, activates the autophagic pathway ^20, 21^. Interestingly, inhibition of ISR, resulted in an increase in mtDNA suggesting that the activation of UPR^mt^ is linked to the enhanced mitochondrial clearance in our model.

Interestingly, mitochondrial stress response and mitophagy transcripts have been found to be upregulated in mild cognitive impairment as well as in mild and moderate AD patients, whereas defective mitophagy may play a role in the disease progression at later stages ^29, 41^. Fang et al have recently shown that the enhancement of mitophagy is able to rescue AD-related pathology in different AD model systems ^29^. In line with this finding, we show that stimulating mitophagy with NMN, a NAD+ booster, significantly improves mitochondrial clearance, with a reduction of Aβ42/Aβ40 ratio and tau hyperphosphorylation. On the contrary, inhibition of UPR^mt^ with ISRIB led to decreased mitochondrial clearance and aggravation of Aβ and tau pathology. Thus, our data suggest a protective role of mitophagy against mitochondrial proteotoxicity induced by PITRM1 deficiency. Interestingly, NMN-related induction of mitophagy was evident only in PITRM1^−/−^ cerebral organoids, while induction of mitochondrial biogenesis was detected in PITRM1^+/+^ organoids upon treatment.

In conclusion, we report a novel cellular model of human PITRM1 deficiency that recapitulates several fundamental pathological aspects of PITRM1-related mitochondrial disease. Using human iPSC-derived cortical neurons and cerebral organoids, we show that loss of PITRM1 function leads to pathological features similar to the ones observed in AD, namely protein aggregation, tau hyperphosphorylation, and neuronal death. We report that PITRM1 deficiency induces impairment of mitochondrial proteostasis and activation of UPR^mt^ that activates cytosolic quality control pathways, such as the UPS and autophagy. The overload of the UPS causes, on the long run, a reduced capacity of degrading cytosolic proteins leading to APP accumulation, increased level of Aβ species, increased Aβ42/40 ratio, and extracellular protein aggregation. Furthermore, we show that, similar to what has been described in AD, enhancing UPR^mt^ and mitophagy ameliorates neuropathological features in primary mitochondrial disease-related neurodegeneration. Importantly, although PITRM1 mutations are relatively rare, the disease mechanisms described in the present study may apply to both primary mitochondrial diseases and more common adult-onset neurological diseases. Thus, our data support a mechanistic link between primary mitochondrial disorders and common neurodegenerative proteinopathies.

## Acknowledgements

We acknowledge the funding support of CoEN Pathfinder II (Ref. 3038, to M.D., M.Z.), the Helmholtz Association (to M.D.), DAAD (PKZ 91723383, to M.J.P.), and Fondazione Umberto Veronesi 2018-2019 (to D.B.) for this project.

## Author contributions

M.D., D.I. and M.J.P. conceived the study. M.D., D.I., M.J.P., V.P., D.B., S.A.K., M.J., M.Z., C.V.contributed to experimental design. D.I. performed gene editing. M.J.P., V.P., G.D.N. performed the most of the experiments with iPSCs and cerebral organoids., R.S performed Meso Scale assay, M.D., D.I., M.J.P., V.P. analysed data. M.D. wrote the manuscript with input and approval from all the authors.

## Conflict of interest

The authors declare no competing interests

## References

1. Johri A, Beal MF. Mitochondrial dysfunction in neurodegenerative diseases. J Pharmacol Exp Ther 2012; 342(3): 619–630.

2. Brunetti D, Torsvik J, Dallabona C, Teixeira P, Sztromwasser P, Fernandez-Vizarra E et al. Defective PITRM1 mitochondrial peptidase is associated with Abeta amyloidotic neurodegeneration. EMBO Mol Med 2016; 8(3): 176–190.

3. Langer Y, Aran A, Gulsuner S, Abu Libdeh B, Renbaum P, Brunetti D et al. Mitochondrial PITRM1 peptidase loss-of-function in childhood cerebellar atrophy. J Med Genet 2018; 55(9): 599–606.

4. Town L, McGlinn E, Fiorenza S, Metzis V, Butterfield NC, Richman JM et al. The metalloendopeptidase gene Pitrm1 is regulated by hedgehog signaling in the developing mouse limb and is expressed in muscle progenitors. Dev Dyn 2009; 238(12): 3175–3184.

5. Stahl A, Nilsson S, Lundberg P, Bhushan S, Biverstahl H, Moberg P et al. Two novel targeting peptide degrading proteases, PrePs, in mitochondria and chloroplasts, so similar and still different. J Mol Biol 2005; 349(4): 847–860.

6. van ’t Hof R, Demel RA, Keegstra K, de Kruijff B. Lipid-peptide interactions between fragments of the transit peptide of ribulose-1,5-bisphosphate carboxylase/oxygenase and chloroplast membrane lipids. FEBS Lett 1991; 291(2): 350–354.

7. Zardeneta G, Horowitz PM. Analysis of the perturbation of phospholipid model membranes by rhodanese and its presequence. The Journal of biological chemistry 1992; 267(34): 24193–24198.

8. Mossmann D, Vogtle FN, Taskin AA, Teixeira PF, Ring J, Burkhart JM et al. Amyloid-beta peptide induces mitochondrial dysfunction by inhibition of preprotein maturation. Cell Metab 2014; 20(4): 662–669.

9. Falkevall A, Alikhani N, Bhushan S, Pavlov PF, Busch K, Johnson KA et al. Degradation of the amyloid beta-protein by the novel mitochondrial peptidasome, PreP. The Journal of biological chemistry 2006; 281(39): 29096–29104.

10. Teixeira PF, Pinho CM, Branca RM, Lehtio J, Levine RL, Glaser E. In vitro oxidative inactivation of human presequence protease (hPreP). Free Radic Biol Med 2012; 53(11): 2188–2195.

11. Pinho CM, Teixeira PF, Glaser E. Mitochondrial import and degradation of amyloid-beta peptide. Biochimica et biophysica acta 2014; 1837(7): 1069–1074.

12. Reinhardt P, Schmid B, Burbulla LF, Schondorf DC, Wagner L, Glatza M et al. Genetic correction of a LRRK2 mutation in human iPSCs links parkinsonian neurodegeneration to ERK-dependent changes in gene expression. Cell stem cell 2013; 12(3): 354–367.

13. Brennand KJ, Simone A, Jou J, Gelboin-Burkhart C, Tran N, Sangar S et al. Modelling schizophrenia using human induced pluripotent stem cells. Nature 2011; 473(7346): 221–225.

14. Lancaster MA, Renner M, Martin CA, Wenzel D, Bicknell LS, Hurles ME et al. Cerebral organoids model human brain development and microcephaly. Nature 2013; 501(7467): 373–379.

15. Branda SS, Cavadini P, Adamec J, Kalousek F, Taroni F, Isaya G. Yeast and human frataxin are processed to mature form in two sequential steps by the mitochondrial processing peptidase. The Journal of biological chemistry 1999; 274(32): 22763–22769.

16. Pellegrino MW, Haynes CM. Mitophagy and the mitochondrial unfolded protein response in neurodegeneration and bacterial infection. BMC Biol 2015; 13: 22.

17. Sorrentino V, Romani M, Mouchiroud L, Beck JS, Zhang H, D’Amico D et al. Enhancing mitochondrial proteostasis reduces amyloid-beta proteotoxicity. Nature 2017; 552(7684): 187–193.

18. Anderson CJ, Bredvik K, Burstein SR, Davis C, Meadows SM, Dash J et al. ALS/FTD mutant CHCHD10 mice reveal a tissue-specific toxic gain-of-function and mitochondrial stress response. Acta neuropathologica 2019; 138(1): 103–121.

19. Forsstrom S, Jackson CB, Carroll CJ, Kuronen M, Pirinen E, Pradhan S et al. Fibroblast Growth Factor 21 Drives Dynamics of Local and Systemic Stress Responses in Mitochondrial Myopathy with mtDNA Deletions. Cell Metab 2019.

20. B’Chir W, Maurin AC, Carraro V, Averous J, Jousse C, Muranishi Y et al. The eIF2alpha/ATF4 pathway is essential for stress-induced autophagy gene expression. Nucleic Acids Res 2013; 41(16): 7683–7699.

21. Rzymski T, Milani M, Pike L, Buffa F, Mellor HR, Winchester L et al. Regulation of autophagy by ATF4 in response to severe hypoxia. Oncogene 2010; 29(31): 4424–4435.

22. Grumati P, Dikic I. Ubiquitin signaling and autophagy. The Journal of biological chemistry 2018; 293(15): 5404–5413.

23. Segref A, Kevei E, Pokrzywa W, Schmeisser K, Mansfeld J, Livnat-Levanon N et al. Pathogenesis of human mitochondrial diseases is modulated by reduced activity of the ubiquitin/proteasome system. Cell Metab 2014; 19(4): 642–652.

24. Durieux J, Wolff S, Dillin A. The cell-non-autonomous nature of electron transport chain-mediated longevity. Cell 2011; 144(1): 79–91.

25. Houtkooper RH, Mouchiroud L, Ryu D, Moullan N, Katsyuba E, Knott G et al. Mitonuclear protein imbalance as a conserved longevity mechanism. Nature 2013; 497(7450): 451–457.

26. Merkwirth C, Jovaisaite V, Durieux J, Matilainen O, Jordan SD, Quiros PM et al. Two Conserved Histone Demethylases Regulate Mitochondrial Stress-Induced Longevity. Cell 2016; 165(5): 1209–1223.

27. Borch Jensen M, Qi Y, Riley R, Rabkina L, Jasper H. PGAM5 promotes lasting FoxO activation after developmental mitochondrial stress and extends lifespan in Drosophila. eLife 2017; 6.

28. Sidrauski C, McGeachy AM, Ingolia NT, Walter P. The small molecule ISRIB reverses the effects of eIF2alpha phosphorylation on translation and stress granule assembly. eLife 2015; 4.

29. Fang EF, Hou Y, Palikaras K, Adriaanse BA, Kerr JS, Yang B et al. Mitophagy inhibits amyloid-beta and tau pathology and reverses cognitive deficits in models of Alzheimer’s disease. Nature neuroscience 2019; 22(3): 401–412.

30. Fang EF, Kassahun H, Croteau DL, Scheibye-Knudsen M, Marosi K, Lu H et al. NAD(+) Replenishment Improves Lifespan and Healthspan in Ataxia Telangiectasia Models via Mitophagy and DNA Repair. Cell Metab 2016; 24(4): 566–581.

31. Hansson Petersen CA, Alikhani N, Behbahani H, Wiehager B, Pavlov PF, Alafuzoff I et al. The amyloid beta-peptide is imported into mitochondria via the TOM import machinery and localized to mitochondrial cristae. Proceedings of the National Academy of Sciences of the United States of America 2008; 105(35): 13145–13150.

32. Manczak M, Anekonda TS, Henson E, Park BS, Quinn J, Reddy PH. Mitochondria are a direct site of A beta accumulation in Alzheimer’s disease neurons: implications for free radical generation and oxidative damage in disease progression. Human molecular genetics 2006; 15(9): 1437–1449.

33. Casley CS, Canevari L, Land JM, Clark JB, Sharpe MA. Beta-amyloid inhibits integrated mitochondrial respiration and key enzyme activities. Journal of neurochemistry 2002; 80(1): 91–100.

34. Aleardi AM, Benard G, Augereau O, Malgat M, Talbot JC, Mazat JP et al. Gradual alteration of mitochondrial structure and function by beta-amyloids: importance of membrane viscosity changes, energy deprivation, reactive oxygen species production, and cytochrome c release. J Bioenerg Biomembr 2005; 37(4): 207–225.

35. Livnat-Levanon N, Glickman MH. Ubiquitin-proteasome system and mitochondria - reciprocity. Biochimica et biophysica acta 2011; 1809(2): 80–87.

36. Nargund AM, Pellegrino MW, Fiorese CJ, Baker BM, Haynes CM. Mitochondrial import efficiency of ATFS-1 regulates mitochondrial UPR activation. Science 2012; 337(6094): 587–590.

37. Shpilka T, Haynes CM. The mitochondrial UPR: mechanisms, physiological functions and implications in ageing. Nat Rev Mol Cell Biol 2018; 19(2): 109–120.

38. Khan NA, Nikkanen J, Yatsuga S, Jackson C, Wang L, Pradhan S et al. mTORC1 Regulates Mitochondrial Integrated Stress Response and Mitochondrial Myopathy Progression. Cell Metab 2017; 26(2): 419–428 e415.

39. Beck JS, Mufson EJ, Counts SE. Evidence for Mitochondrial UPR Gene Activation in Familial and Sporadic Alzheimer’s Disease. Curr Alzheimer Res 2016; 13(6): 610–614.

40. Gerakis Y, Hetz C. Brain organoids: a next step for humanized Alzheimer’s disease models? Mol Psychiatry 2019; 24(4): 474–478.

41. Kerr JS, Adriaanse BA, Greig NH, Mattson MP, Cader MZ, Bohr VA et al. Mitophagy and Alzheimer’s Disease: Cellular and Molecular Mechanisms. Trends in neurosciences 2017; 40(3): 151–166.

